# New Insights into Molecular Basis Identification of Three Novel Strains of the Bacillus Subtilis Group Produce Cry Proteins Isolated from Soil Samples in Adana, Turkey

**DOI:** 10.1101/2021.12.24.474129

**Authors:** Semih Yılmaz, Abeer Babiker Idris, Abdurrahman Ayvaz, Aysun Çetin, Funda Ülgen, Mustafa Çetin, Berkay Saraymen, Mohamed A. Hassan

## Abstract

**Aims:** This study aimed to analyze the evolutionary relationship between *Bacillus* species isolated from agricultural soil using in-silico tools.

**Methods and Results:** Across-sectional study was conducted in Adana province, in Turkey. A total of 120 *Bacillus* species were isolated from 80 soil samples. However, the phylogenetic tree diverged into two lineages; one belongs to *B. subtilis* group while the other belongs to *B. cereus* group. Interestingly, three native strains (*SY27.1A, SY35.3A*, and *SY58.5A*), which produce Cry proteins, shared high similarity with *B. subtilis* group (over 99%) and less than 95% similarity with known *B. thuringiensis* and other species of *B. cereus* group. Furthermore, 11 canonical SNPs (canSNPs) were identified in strains that belong to *B. pumilus* group when compared with *B. subtilis* reference sequences.

**Conclusions:** Phylogenetic analysis of *16S rRNA* sequences was found valuable for differentiation between *Bacillus* species isolated from soil samples. In addition, SNPs analysis provided more intra-specific information in the cases of *B. subtilis* group.

**Significance and Impact of Study:** A detailed analysis was provided for the SNPs present in a conserved region of *16S rRNA* gene of *Bacillus* species. Also, we proposed three novel *Bacillus* strains that produce Cry proteins and belong to *B. subtilis* group.

## 1. Introduction

*Bacillus* is agriculturally important insecticidal bacterial genus that naturally inhabit the phyllosphere and rhizosphere (1). It consists of a heterogeneous group of Gram-positive, endospore-forming, aerobic or facultative anaerobic organisms (2). Most members of the genus *Bacillus* have the ability to produce antibiotics, enzymes, vitamins, proteins, or secondary metabolites that are capable to induce defense mechanisms and promote growth in animals and plants (3, 4). Benefiting from their metabolic diversity and spore dispersal, *Bacillus* is ubiquitous in various natural environments especially terrestrial environments (5). At the time of writing, the genus *Bacillus* consisted of more than 408 species with validly published names (LPSN, http://www.bacterio.net), only 54 species of them were reported before 2000 (6).

Analysis of *16S rRNA* gene, the “ultimate molecular chronometer”, has been extensively applied for bacterial phylogeny and taxonomy which resulted eventually in the establishment of large public-domain databases (7-11). The *16S rRNA* gene characterizes by several properties which include being present in all bacteria, thus it is a universal target for bacterial identification and characterization (12, 13). In addition, the function of *16S rRNA* has not changed over a long period, i.e. random sequence changes are more likely to reflect the microbial evolutionary change (phylogeny) (11), and any introduction of selected changes in one domain does not greatly affect sequences in other domains (13, 14). Based on phylogenetic analysis of the *16S rRNA* gene, the species and strains in *Bacillus* are divided into five groups: *B. cereus, B. megaterium, B. subtilis, B. circulans* and *B. brevis* groups (15).

The *B. subtilis* group is a tight assemblage of closely related species which includes *B. subtilis, B. amyloliquefaciens, B. atrophaeus, B. axarquiensis, B. malacitensis, B. mojavensis, B. sonorensis, B. tequilensis, B. vallismortis* and *B. velezensis* (16). These species share high genetic homogeneity (over 99.5%) and cannot differentiate on the basis of phenotypic or biochemical characteristics (15, 16). In addition, *B. pumilus* and their relatives belong to the *subtilis* group (17). The *B. pumilus* group, which is a large group of *Bacillus*, composed of *B. pumilus, B. altitudinis, B. safensis, B. zhangzhouensis, B. xiamenensis*, and *B. australimaris* (5).

The bacteria of *B. cereus* group share high genetic homogeneity despite their phenotypic diversity, with over 97% *16S rRNA* sequence similarity among *B. cereus, B. anthracis, B. thuringiensis, B. weihenstephanensis, B. mycoides, B. pseudomycoides, B. cytotoxicus, B. gaemokensis* and *B. manliponensis* (18). Moreover, this group is of interest to researchers, especially *B. thuringiensis*, because of their significance in agriculture, industry and medicine (19). *B. thuringiensis* acts as a biological control agent against different phytopathogenic organisms due to their ability to produce insecticidal proteins (Cry, Vip, Sip, Bin, etc), fungicidal metabolites (iturin, fengycin, surfactin, zwittermycin, etc) (20, 21). Also, *B*. *thuringiensis* can promote plant growth by producing ACC deaminase, phosphatases, siderophore, etc (22, 23). However, the isolation and characterization of native *Bacillus* species or strains, especially from agricultural soil, should receive a good attention because of their wide potential biological products with immense applications. In addition, there are several studies on the isolation and characterization of native *Bacillus* strains of soil in order to identify novel toxins with high level of toxicity and effective against agricultural pests (24-26). Finding of novel toxins produce from *Bacillus* species, especially *B. thuringiensis*, will delay the resistance within the pests due to the use of existing *Bacillus* toxins. Therefore, in this study, we aimed to characterize and establish a phylogenetic relationship between *Bacillus* species isolated from agricultural soil in Adana, one of the most fertile agricultural area in Turkey, by reconstructing *16S rRNA* phylogenetic trees using in silico tools. Also, a detailed analysis was provided, for the SNPs present in a conserved region of *16S rRNA* gene of *B. cereus* group, *B. Subtilis* group and *B. pumilus* group. The results of canonical SNPs (canSNPs) are of great significance for the design of primers or probes specific to a strain, species, or group of species.

## 2. Materials and Methods

### 2.1 Study Settings and Sample collection

A cross-sectional study was conducted in Adana province, which is located in the southern region of Turkey. Adana province is divided into 13 districts with different texture. For the isolation of bacterial strains, 80 different soil samples were collected throughout Adana province from different altitudes ranging from 0 to 1582 meters. Soil samples have been taken in a depth of 2-10 centimeters and stored in sterile tubes at 4°C for the studies.

### 2.2 Bacterial Isolation and Identification

Isolation processes has been performed according to the method of Travers *et al*. (27). One gram of soil sample was inoculated in LB medium (pH 6.8±2) including 0.25M sodium acetate in a shaking incubator at 200 rpm at 30°C for 4 hours. After the incubation step, 1.5 ml of liquid samples have been transferred to a sterile Eppendorf tube and exposed to 80°C for 10 minutes to kill the vegetative bacterial forms. A 20-50 ul of samples were spread on LB agar plates and incubated overnight at 30°C. The colonies with morphological differences have been spread on agar plates and pure colonies were obtained. Pure colonies were incubated in 5ml LB broth (pH 6.8±2) in 50 ml tubes at 200 rpm and 30°C overnight.

Then the colonies were homogenized in 400 μl of sterile dH_2_O in microfuge tubes and 10 ul were added onto sterilized Watman no:1 paper disc with 0.4 mm diameter. The discs were then placed into Potato Dextrose agar (PDA) plates and incubated overnight at 30°C. Morphologically pale-yellow, grayish white, pale-pink, ciliated, or wrinkled ends, round-shaped outlines were selected for Gram stain (28). Gram-positive colonies were further investigated for spore production using Malachite green (5g /100ml) staining as previously described (29).

### 2.3 Characterization of para-sporal inclusions

To characterize para-sporal inclusions, the bacteria were incubated in 150 ml of 3T medium (2 g triptose, 3 g triptone, 1.5 g yeast extract, 6 g NaH_2_PO_4_, 0.005 g MnCl2 and 7.1 g Na_2_HPO_4_) at 200 rpm and 30 °C for 7 days to induce sporulation (27). Then, to harvest spore-crystal mixtures, the suspensions were centrifuged at 15000 ×g and 4 °C for 10 min. After that, the mixtures were suspended in dH_2_O on microscope slides and fixed. Finally, the slides were sputter coated with 10 nm Au/Pd using a SC7620 Mini-sputter coater and viewed using a LEO_4_40 scanning electron microscope (SEM) at 20kV beam current (30, 31) in this study reference standard *B. thuringiensis* strains such as *Bt. kurstaki HD1, Bt. kurstaki HD73, Bt. aizawai* (Universidad Nacional Autonoma de Mexico Biyotechnology Institute), *Bt. morrisoni, Bt. israelensis* (Pasteur Institute, Paris, France) and, *Bt. tenebrionis* (Plant Genetic Systems, J. Plateaustroat 22, 900 Gent, Belgium) were used for comparison with the native local isolates.

### 2.4 Genetics analysis

#### 2.4.1 Determination of the insecticidal *crystal* genes (*cry*) carrying isolates

Extraction of DNA was performed according to a previously described method (32, 33). Briefly, the bacteria were grown in LB medium for overnight, and then a loopful of cells was placed into 400 μl sterile dH_2_O. Then the mixture was boiled for 10 min to lyse the cells. The resulting cell lysate was centrifuged for 10 sec at 10.000 rpm and the supernatant was used as DNA templates for PCR reactions. The extracted DNA from the isolates was used to determine the *cry* genes carrying strains. In our previous work, we have characterized the isolates using *cry* genes *cry1Ab/ Ac, cry1Aa/Ad, cry2, cry5, and cry9C, cry1C, cry1Ad, cry1Ac, cry1D, cry1B, cry3-7-8, cry4A, cry9A*, and *cry11A/B* (30, 31).

#### 2.4.2 Amplification and sequencing of *16S rRNA* gene

The extracted DNA was used to amplify the *16S rRNA* gene of Bacillus species using universal primers with the following sequences: F: 5’-AAA CTY AAA KGA ATT GAC GG-3’ and R: 5’-ACG GGC GGT GTG TRC-3’. The thermal procedures were performed with ABI veriti Thermocycler and the PCR mixtures contained 2.3 mM MgCl2, 1x Taq buffer, 0.2 mM dNTP mix, 0.3 pmol for each primer, 0.5 U Taq DNA polymerase, and 30–100 ng template DNA. The PCR conditions were an initial denaturation 94°C 5 min, then 40 cycles of denaturation at 94°C for 1min, primer annealing at 48°C for 1 min, extension at 72°C for 2min, and then additional extension step 72°C for 10min. The size of expected PCR products was 850 bp.

For sequencing, the DNA fragments were extracted from the gel using a EasyPure® Quick Gel Extraction Kit (EG101-01) according to the manufacturer’s instruction. Then the PCR products of 21 samples, which have the clearest bands, were sent for commercial DNA purification and Sanger dideoxy sequencing by DETAGEN Genetic Diagnostics Center Inc., Turkey.

### 2.5 Bioinformatics analysis

#### 2.5.1 Sequence and SNP analysis

The two chromatograms for each strain (forward and reverse) were visualized, checked the quality, and analyzed using the Finch TV program version 1.4.0 (34). The bacterial strains were identified by searching for their homology among published reference sequences using the nucleotide Basic Local Alignment Search Tool (BLASTn; https://blast.ncbi.nlm.nih.gov/) (35). To determine the SNPs, multiple sequences alignment (MSA) was accomplished with reference sequences of *Bacillus* species using BioEdit software (36) and MEGA version 7.0 software (37). This MSA facilitated the use of polymorphisms to detect potential relationships between the *Bacillus* strains and species. In addition, the detected SNPs were carefully reviewed by eye using the Finch TV software; and polymorphisms present in both the forward and reverse strands were considered.

#### 2.5.2 Molecular phylogenetic analysis

For building the phylogenetic tree, the studied sequences and their highly similar references sequences that retrieved from the NCBI GenBank were subjected to Gblocks software to eliminate poorly aligned positions and divergent regions of aligned sequences, so the alignment becomes more suitable for phylogenetic analysis (38). The molecular evolutionary analyses were conducted with MEGA 7.0 software (37) using the maximum likelihood (ML) method and neighbour-joining (NJ) method (39, 40). The Kimura 2-parameter (K2+G) model from the substitution (ML) model was used with 1000 bootstrap replicates to construct distance-based trees (41, 42).

## 3. Results

### 3.1 Bacterial isolation and characterization

A total of 120 *Bacillus* species were isolated from 80 soil samples. Eighty-eight of *Bacillus* isolates harbored *cry* genes. The *cry* genes were determined and characterized using conventional PCR. Also, spore–crystals of some of the samples were examined under the SEM. The results in details were presented in our previous works (30, 31). In the current study, *16S rRNA* of 21 *Bacillus* species were amplified and sequenced to construct a phylogenetic tree. Among them, six strains were characterized by producing Cry proteins. The morphology of spore–crystals and *cry* genes profiles of the native *Bacillus* isolates are illustrated in Table 1. The nucleotide sequences of the *16S rRNA* were deposited in the GenBank database under the following accession numbers: from OK428682 to OK428687 and from OK384678 to OK384692.

**Table 1.**
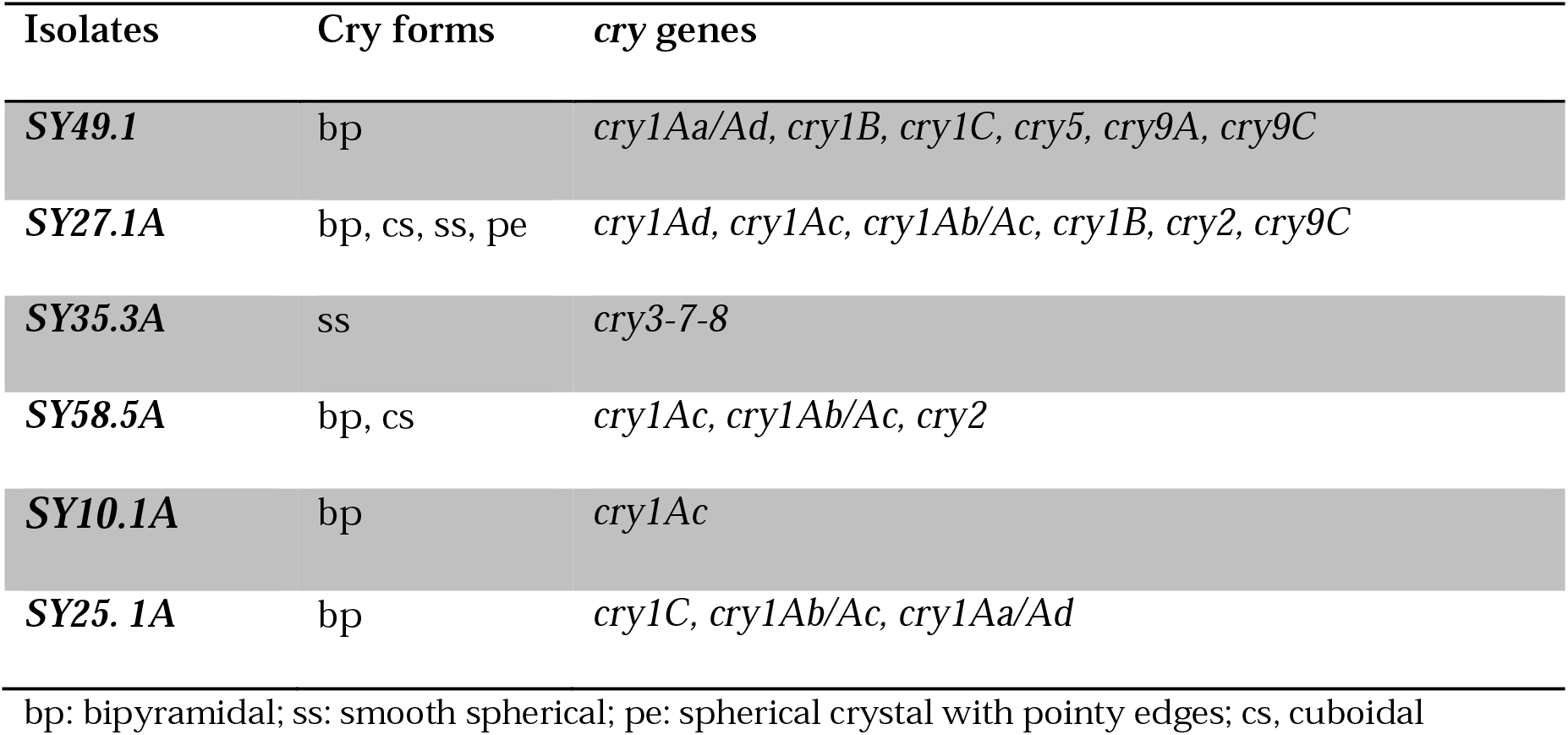
The morphology of spore–crystals and *cry* gene profiles of the native *Bacillus* isolates

| Isolates | Cry forms | *cry* genes |
| --- | --- | --- |
| **SY49.1** | bp | *cry1Aa/Ad, cry1B, cry1C, cry5, cry9A, cry9C* |
| **SY27.1A** | bp, cs, ss, pe | *cry1Ad, cry1Ac, cry1Ab/Ac, cry1B, cry2, cry9C* |
| **SY35.3A** | ss | *cry3-7-8* |
| **SY58.5A** | bp, cs | *cry1Ac, cry1Ab/Ac, cry2* |
| **SY10.1A** | bp | *cry1Ac* |
| **SY25. 1A** | bp | *cry1C, cry1Ab/Ac, cry1Aa/Ad* |

bp: bipyramidal; ss: smooth spherical; pe: spherical crystal with pointy edges; cs, cuboidal

### 3.2 Sequencing analysis of *16S rRNA* gene

Twenty-one isolates of *Bacillus* species were subjected to PCR amplification and nucleotide sequencing using universal *16S rRNA* primers but specific for a conserved region of the gene which was located between 629 bp and 1552 bp. The sequences of the studied strains were aligned with reference sequences of *Bacillus* species retrieved from NCBI databases. The information about the retrieved strains is given in supplementary Table S1.

As presented in Figure 1, 11 strains revealed high similarity with the *B. cereus* group (over 99%), see Figure 1. Among them, seven strains, which were found to be homogenous, revealed two nucleotide variations (A1015C and A1146T). Numbers are given in all sequences in accordance with numbering in the *B. thuringiensis* genome (NZ_CM000747). Six of these strains (*SY49.1, SY10.1A*, and *SY25.1A*) were characterized by the production of Cry proteins. While six and four of the studied strains exhibited high similarities to *B. subtilis* group and *B. pumilus* group, respectively (Figure 3). Interestingly, three of native strains (*SY27.1A, SY35.3A, SY58.5A*), which produce Cry proteins, shared high similarity with *B. subtilis* group (over 99%) and less than 95% similarity with known *B. thuringiensis* and other species of *B. cereus* group, see Figure 2.

**Figure 1.**
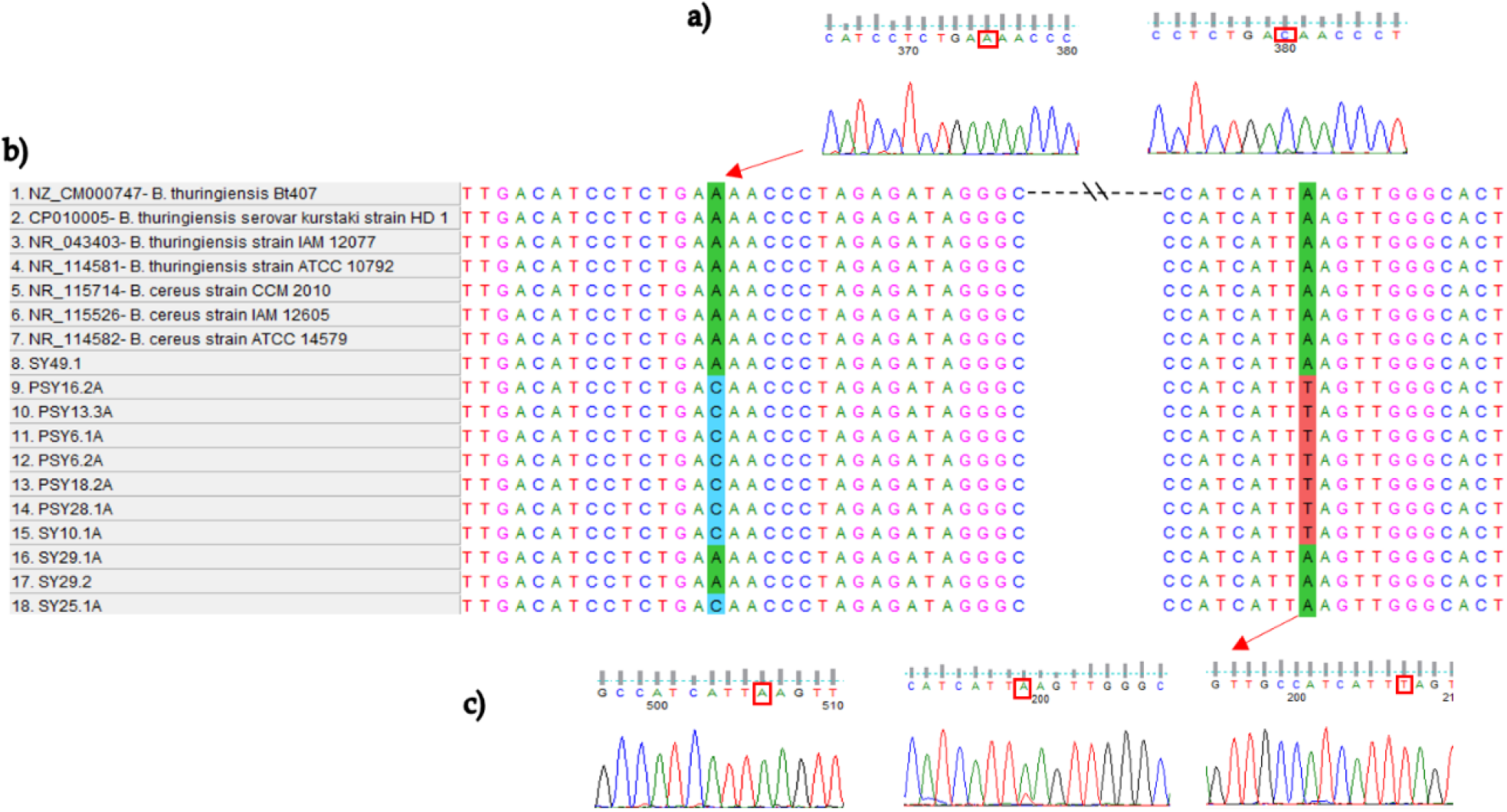
1a) and 1b) Sequencing results of chromatograms using Finch TV software show nucleotide changes in the *16S rRNA* gene of *B. cereus* / *B. thuringiesis* strains illustrated by squares. 1c) Multiple sequences alignment of the native *Bacillus* strains that belong to *B. cereus* group using ClustalW.

**Figure 2.**
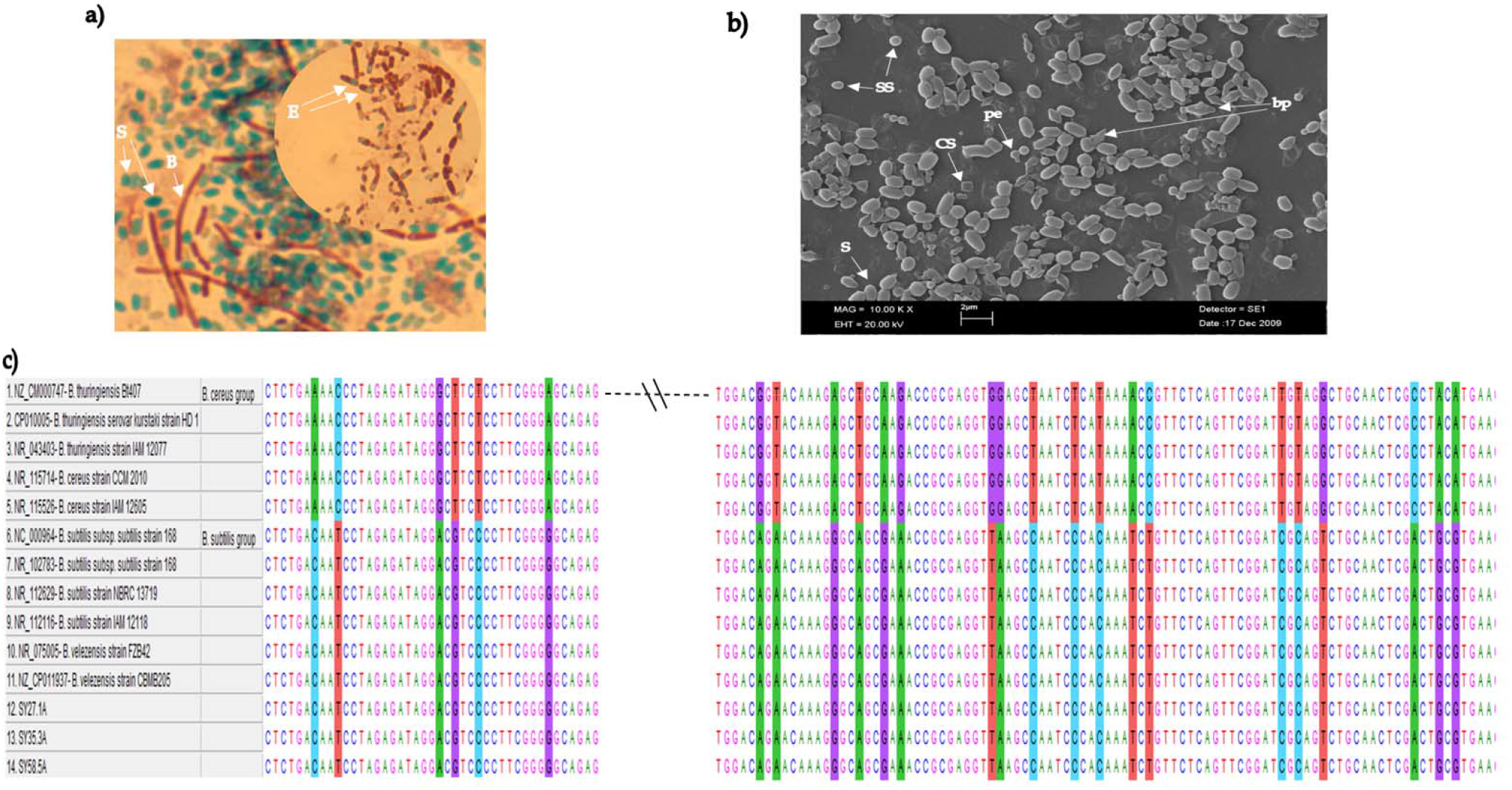
2a) Microscopic view of native *Bacillus* strains after spore staining (1000x). B, *Bacillus* bacteria; S, spores; E, endospores. 2b) Scanning electron microscopy (SEM) image of the different types of crystal morphologies and the spores (S) produced by native *Bacillus* strain *SY27.1A* isolated from soil sample. SS, smooth spherical crystal; pe, spherical crystal with pointy edges; bp, bipyramidal crystal; cs, cuboidal crystal. 2c) Multiple Sequence Alignment (MSA) of *16S rRNA* sequences of the three native strains (*SY27.1A, SY35.3A, SY58.5A*) shows high similarity with *B. subtilis* group rather than species of *B. cereus* group.

**Figure 3.**
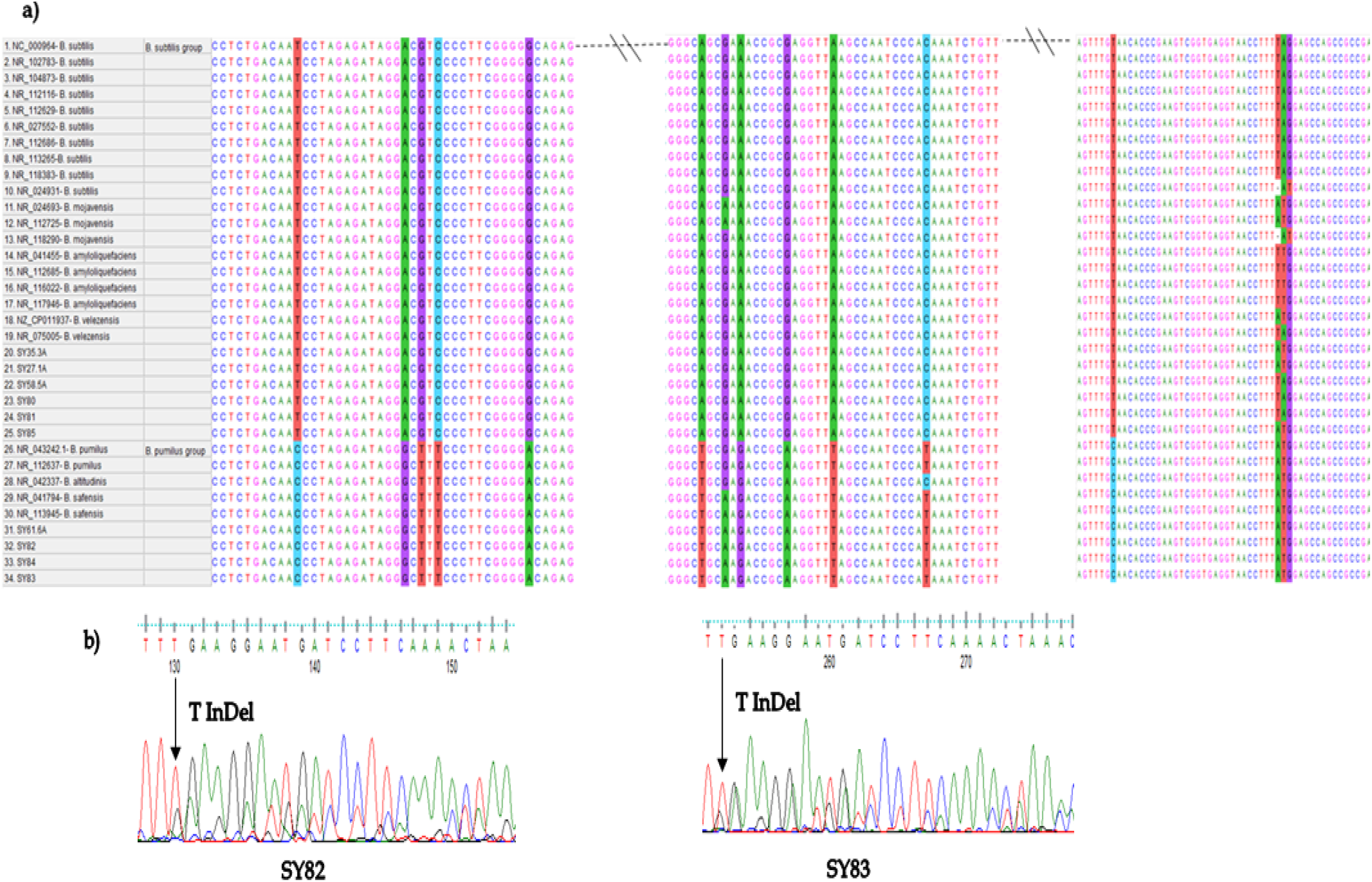
3a) Multiple sequences alignment (MSA) of *B. subtilis* group shows 11 canonical SNPs (canSNPs) in strains that belong to *B. pumilus* groups when compared with *B. subtilis* reference sequences. 3b) Sequencing results of chromatograms illustrate the loss of synchronicity in *SY82* and *SY83* strains due to a nucleotide deletion or insertion within one or more of the *16S rRNA* genes among the multiple *rRNA* operons in the genome.

### 3.3 Molecular phylogenetic analysis

The evolutionary analysis of 21 native Turkish *Bacillus* strains, based on *16S rRNA* gene, was conducted with reference sequences of *Bacillus* species retrieved from NCBI GenBank databases. Neighbor-joining (NJ), maximum-likelihood (ML) analyses were performed with MEGA 7 (37). The topology of the ML and NJ trees was similar, and the bootstrap supports of the NJ tree were approximately higher than those of ML. The phylogenetic tree diverged into two lineages; one belongs to *B. subtilis* group while the other belongs to *B. cereus* group (Figure 4).

**Figure 4.**
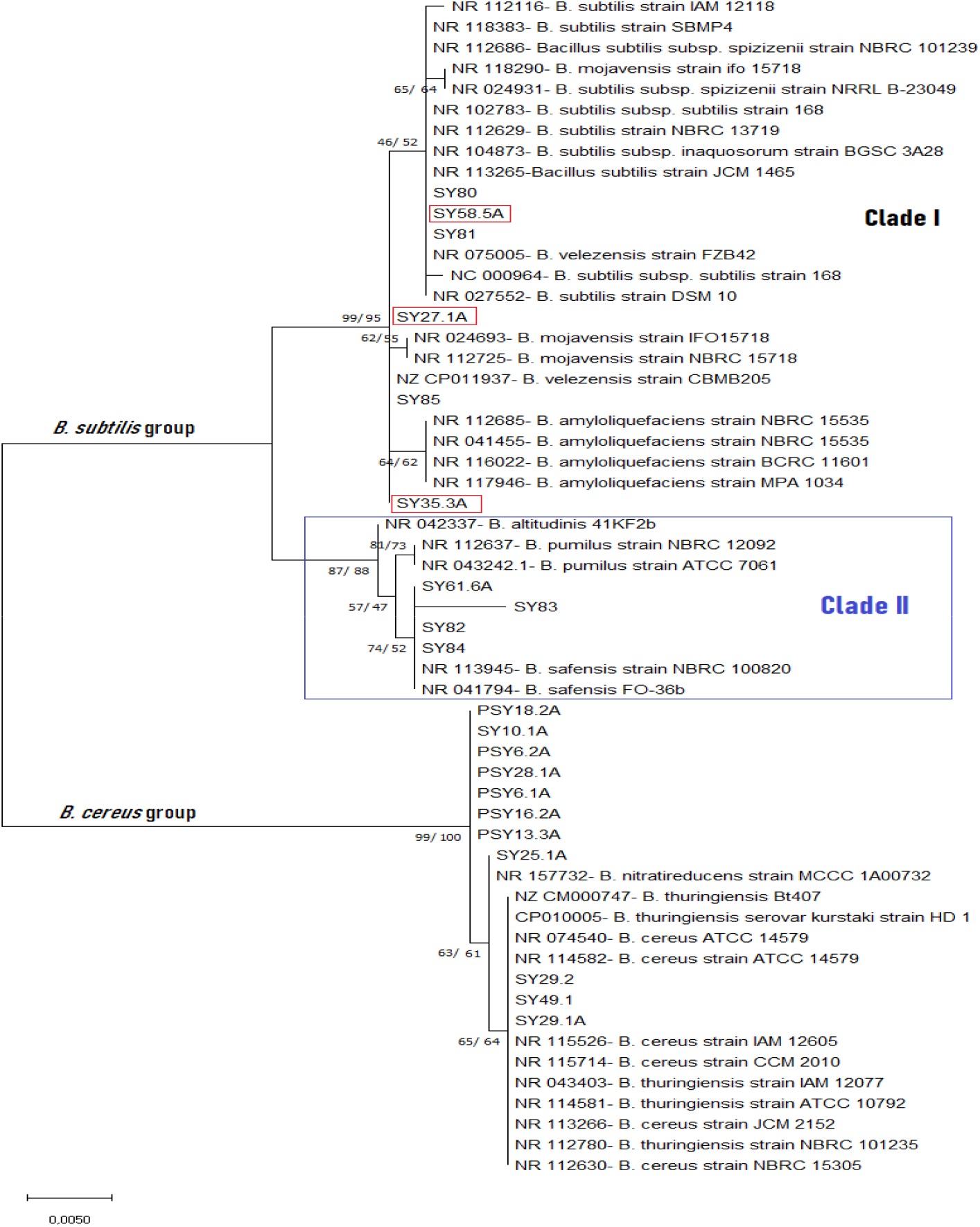
Maximum-Likelihood Phylogenetic tree of native *Bacillus* strains isolated from Turkey. The percentage of replicate trees (1000 replicates) are shown next to the branches from both NJ (before the slash ‘/’) and ML (after the slash ‘/’) analyses. The ML tree is only shown here, because the ML tree was very similar to the NJ tree. The evolutionary distance was computed using (K2+G) model and evolutionary analyses were conducted using MEGA7.

The lineage of *B. subtilis* group branched into two major clades (I and II). In clade I, six strains (SY27.1A, SY35.3A, SY58.5A, SY80, SY81 and SY85) were clustered with *B. subtilis, B. amyloliquefaciens, B. velezensis* and *B. mojavensis*. However, SY27.1A, SY35.3A and SY85 strains were closely related to *B. velezensis* strain CBMB205 and they shared nucleotide variations at TA1461-1462AT. While in clade II, four strains (*SY61.6, SY82, SY83* and *SY84*) were grouped with *B. pumilus* group which comprises *B. pumilus, B. safensis*, and *B. altitudinis*. In addition, all strains in clade II shared 11 nucleotide variations with *B. pumilus* group (T1017C, A1030G, G1032T, C1034T, G1045A, A1265T, A1270G, G1276A, A1282T, T1432C and A1485G), see Figure 3a. Four mutations (C971G, C1316T, C1330T and A1543C) in strain A made it a separate minor clade. Interestingly, three native strains (*SY27.1A, SY35.3A* and *SY58.5A*) that characterized by producing Cry proteins, which is often considered as a feature of *B. thuringiensis*, were clustered with *B. subtilis* group (Figure 2). The lineage of *B. cereus* group involved 11 strains. Three of them (*SY49.1, SY10.1A*, and *SY25.1A*) were characterized by producing Cry proteins. Moreover, seven strains shared a common ancestor and were characterized by two nucleotide variations (A1015C and A1146T). However, strain *SY25.1A* and *B. nitratireducens* strain *MCCC 1A00732* were sisters with a bootstrap value of 63%, see Figure 4 for more illustration.

## 4. Discussion

In this study, we observed three novel Gram-positive bacilli (*SY27.1A, SY35.3A*, and *SY58.5A*) which produce Cry proteins but, based on the analysis of *16S rRNA* gene, they are unlikely to belong to the known *B. thuringiensis* or other species of *Bacillus cereus* group. Although the defining feature of the *B. thuringiensis* species is the ability to express Cry proteins (20), the analysis of *16S rRNA* gene sequences of these strains showed that they were sharing over 97% similarity with *B. subtilis* group and less than 95% similarity with the known *B. thuringiensis* and other species of *B. cereus* group. However, in prokaryotes taxonomy, *16S rRNA* gene sequence identity of 97 % is generally used as a threshold value for species definition, therefore, strains with less than 97.5% identity are unlikely to be related at the species level (43, 44). This finding is partially in agreement with a previous study that systematically searched for Cry proteins expressed by *Bacillus* species, other than *B. thuringiensis*, genomes using conserved sequences from the C-terminal half of reported Cry proteins in hidden Markov Model (HMM) profiles (45). Interestingly, there were 174 Cry protein sequences were observed, as expected most of them were in *B. thuringiensis* genomes, but 42 were found in other species. In addition, several studies reported the presence of Cry proteins in other Bacilli, such as *P. popilliae, C. bifermentans, L. sphaericus* and *P. lentimorbus* (46-49). Nevertheless, the great diversity of Cry proteins may indicate that this family of proteins may not be restricted only to the *B. thuringiensis* species (20), and their dispersion and role in nature are could be much wider. Therefore, further studies, either based on an in-silico procedure or the use of a large data collection of different species of bacilli, are recommended to search for new Cry proteins with higher toxicity or different mode of action, which may render alternatives in case of resistance development. However, the development of resistance to insecticidal *B. thuringiensis* proteins has been documented which raise concerns about the adequacy of current resistance management strategies (54). Hence, continuous searching for novel *Bacillus* species with novel insecticidal genes to delay the development of insect resistance is of the utmost importance. In this connection, to isolated novel *Bacillus* strains, we collected soil samples throughout Adana province. This region is rich in biodiversity due to its unique climate and geographical location that is situated on the fertile and watery delta of Seyhan and Ceyhan rivers, furthermore in this study, the genetic identity based on *16S rRNA* sequences indicated that three native strains of *Bacillus* (SY27.1A, SY35.3A, and SY58.5A), which produce Cry proteins were close relatives of the *B. subtilis* group and appeared to be discrete from the *B. cereus* group. However, according to the low discrimination of the *16S rRNA* gene between *Bacillus* species, it cannot be assigned accurately as a certain species (55), therefore, complete genome sequencing of these bacterial strains is recommended. While, the other ten *Bacillus* strains, which clustered with *B. cereus* group, showed diversity into two minor clades. Seven of them shared a common ancestor and were characterized by two nucleotide variations (A1015C and A1146T). This finding is in agreement with other studies conducted in different countries which showed diversity in *B. cereus* / *B. thuringiesis* strains isolated from soil and other natural sources (24, 26, 56-58).

Furthermore, in the phylogenetic tree of the native strains with global reference sequences of *Bacillus* species, the lineage of *B. subtilis* group were branched into two major clades. Clade I contained *B. amyloliquefaciens, B. velezensis, B. mojavensis* along with *B. subtilis* which is not entirely unexpected since these species share a remarkably high level of *16S rRNA* gene sequence similarity to *B. subtilis* (often 99 % or greater) (16). While in clade II, the members of *B. pumilus* group (*B. pumilus, B. altitudinis*, and *B. safensis*) were clustered together along with four strains (*SY61.6A, SY82, SY83*, and *SY84*). They shared 11 nucleotide variations when their *16S rRNA* sequences compared with *B. subtilis* reference sequences, see Figure 3a. In 1973, Gordon *et al*. speculated that *B. pumilus* might one day be considered a variety of *B. subtilis* rather than a separate species once more data were collected (59). Intriguingly, our findings are in agreement with Rooney *et al*. results which clearly indicated that the *B. pumilus* forms a clade distinct from *B. subtilis* (16).

However, single nucleotide polymorphism (SNP) analysis has emerged as one of the most useful molecular methods proposed for microbial characterization and improving discrimination among closely related species (60-62). Accordingly, we used universal primers of *16S rRNA* in order to produce a mixture of amplicons from all *rRNA* operons in the genome. Although there are small differences exhibit by the multiple *rRNA* gene copies in each genome, these differences do not invalidate bacterial identification and characterization based on *16S rRNA* sequences (63, 64). Moreover, these differences between *rRNA* operons appear in multiple peaks (two or more) at a single nucleotide position in the case of SNPs. But in the case of InDels variations, the sequence loses synchronicity and makes an abrupt change, from clean to dirty, following an InDel mutation (65), for more illustration see Figure3b. In the present study, we provided a detailed analysis of the SNPs present in the *16S rRNA* gene of *Bacillus* species isolated from soil samples. Of great significance, dual peaks (A and T) at position 1146, which were previously reported to be specific to *B. anthracis* (66), were detected in three *B. thuringiensis /B. cereus* strains (*PSY6.1A, PSY6.2A* and *SY10.1A*). But in all other strains, a single peak (either A or T) was detected. This finding is in accordance with a study conducted by Hakovirta *et al*. which found five *B. cereus* strains and three *B. thuringiensis* strains also had A and T peaks (65). Hence, the SNP at nucleotide position 1146 is not necessarily reliable for identifying *B. anthracis*. While, in this study, all *B. thuringiensis* had only G at position 1139 which is another SNP proposed by Hakovirta *et al*. to be unique to *B. anthracis* (65), and it appears to be more reliable for identifying *B. anthracis*. Regarding the *B. subtilis* group, we identified 11 canonical SNPs (canSNPs) in strains that belong to *B. pumilus* groups (clade II) when compared with *B. subtilis* reference sequences. Canonical SNPs (canSNPs) are useful and diagnostic SNPs that used for identifying long branches or key phylogenetic positions (67). In addition, G1268A and C1294T SNPs were specific to *B. safensis* and *B. altitude*, respectively. Also, strain *SY83* characterized by four nucleotide variations SNPs (C971G, C1316T, C1330T and A1543C). These findings are partially in agreement with a study conducted by Moorhead *et al*. in which strains of *Listeria monocytogenes* were partitioned into three previously described clonal lineages using a phylogenetic approach to detect a small number of SNPs in the *sigB* gene (68). However, a number of researchers have found that a small number of SNPs can be used to effectively identify genetic groups (61, 67, 69). Also, Keim *et. al*. proposed canSNPs to define the *B. anthracis* lineage that contains the Ames strain (67). The limitations of the present study include the relatively small sample size and the phylogenetic tree was built based on the *16S rRNA* gene only. Hence, further studies with large sample size and molecular techniques, such as multilocus sequence analysis (MLST) and whole genome sequencing (WGS), that are used to differentiate the closely related microbial species like *Bacillus* species are recommended. However, in this work, the SNPs analysis provided more intra-specific information than phylogenetic analysis in the cases of *B. subtilis* group. Eleven canSNPs were identified in strains that belong to *B. pumilus* groups when compared with *B. subtilis* reference sequences. In addition, these canSNPs in the conserved region of *16S rRNA* gene may provide important information for the design of primers and probes for Real-Time PCR, multiplex-PCR and microarray systems which is widely used for detection and typing purposes.

In conclusion, the phylogenetic tree diverged into two lineages; one belongs to *B. subtilis* group while the other belongs to *B. cereus* group. Interestingly, three of native strains (*SY27.1A, SY35.3A*, and *SY58.5A*), which produce Cry proteins, shared high similarity with *B. subtilis* group (over 99%). An 11 canSNPs were identified in strains that belong to *B. pumilus* groups when compared with *B. subtilis* reference sequences. These canSNPs in the conserved region of *16S rRNA* gene may provide important information for the design of primers and probes which is widely used for detection and typing purposes.

## Supplementary file

Table S1. Information of the reference sequences of *Bacillus* species that were retrieved from NCBI databases.

## Acknowledgment

We gratefully acknowledge the Genome and stem cell (GenKök) research center for their cooperation and supporting the study experiments.

## Authors’ contributions

**Semih Yılmaz:** Conceptualization, funding acquisition, methodology, writing - review & editing, and supervision; **Abeer Babiker Idris**: conceptualization, methodology, investigation, software, data curation and formal analysis, writing - original draft; and writing - review & editing; **Abdurrahman Ayvaz:** funding acquisition, methodology and supervision; **Aysun Çetin:** methodology; **Funda Ülgen:** methodology and writing - original draft; **Mustafa Çetin:** funding acquisition, and supervision; **Berkay Saraymen:** methodology and data curation and formal analysis; **Mohamed A. Hassan:** conceptualization, methodology, writing - review & editing, and supervision.

## Availability of data

All data generated or analyzed during this study are included in the manuscript.

## Ethical Approval

This study was approved by Erciyes University, Faculty of Agriculture, Department of Agricultural Biotechnology.

## Consent of participants

Not applicable

## Highlights

- Isolation and characterization of native *Bacillus* strains from agricultural soil should receive a good attention because of their wide potential biological products with immense applications.
- Identification of three native strains (*SY27.1A, SY35.3A*, and *SY58.5A*), which produce Cry proteins, shared high similarity with *B. subtilis* group (over 99%) and less than 95% similarity with known *B. thuringiensis* and other species of *B. cereus* group.
- Eleven canonical SNPs (canSNPs) were detected in strains that belong to *B. pumilus* group when compared with *B. subtilis* reference sequences.
- The SNP at nucleotide position 1146 is not necessarily reliable for identifying *B. anthracis*, while G at position 1139, which is another SNP proposed to be unique to *B. anthracis*, appears to be more reliable for identifying *B. anthracis*.
- Phylogenetic analysis of *16S rRNA* sequences was found valuable for differentiation between *Bacillus* species isolated from soil samples.

