## supplemental Table S1 for "New Insights into Molecular Basis Identification of Three Novel Strains of the Bacillus Subtilis Group Produce Cry Proteins Isolated from Soil Samples in Adana, Turkey"

**Table S1.** Information of the reference sequences of *Bacillus* species that were retrieved from NCBI databases.

| Name of strains | *Bacillus* species | Accession No. | Country | Date of submission |
| --- | --- | --- | --- | --- |
| *SY27.1A* | *B. subtilis group* | OK428682 | Turkey | 13- Oct -2021 |
| *SY81* | *B. subtilis group* | OK428683 | Turkey | 13- Oct -2021 |
| *SY35.3A* | *B. subtilis group* | OK428684 | Turkey | 13- Oct -2021 |
| *SY82* | *B. pumilus group* | OK428685 | Turkey | 13- Oct -2021 |
| *SY83* | *B. pumilus group* | OK428686 | Turkey | 13- Oct -2021 |
| *SY85* | *B. subtilis group* | OK428687 | Turkey | 13- Oct -2021 |
| *SY29.2* | *B. cereus group* | OK384678 | Turkey | 12- Oct -2021 |
| *SY49.1* | *B. thuringiensis* | OK384679 | Turkey | 12- Oct -2021 |
| *PSY16.2A* | *B. cereus group* | OK384680 | Turkey | 12- Oct -2021 |
| *PSY13.3A* | *B. cereus group* | OK384681 | Turkey | 12- Oct -2021 |
| *PSY6.1A* | *B. cereus group* | OK384682 | Turkey | 12- Oct -2021 |
| *PSY28.1A* | *B. cereus group* | OK384683 | Turkey | 12- Oct -2021 |
| *PSY6.2A* | *B. cereus group* | OK384684 | Turkey | 12- Oct -2021 |
| *PSY18.2A* | *B. cereus group* | OK384685 | Turkey | 12- Oct -2021 |
| *SY10.1A* | *B. thuringiensis* | OK384686 | Turkey | 12- Oct -2021 |
| *SY25.1A* | *B. nitratireducens* | OK384687 | Turkey | 12- Oct -2021 |
| *SY29.1A* | *B. cereus group* | OK384688 | Turkey | 12- Oct -2021 |
| *SY80* | *B. subtilis group* | OK384689 | Turkey | 12- Oct -2021 |
| *SY58.5A* | *B. subtilis group* | OK384690 | Turkey | 12- Oct -2021 |
| *SY61.6A* | *B. cereus group* | OK384691 | Turkey | 12- Oct -2021 |
| *SY84* | *B. pumilus group* | OK384692 | Turkey | 12- Oct -2021 |
| *strain CCM 2010* | *B. cereus* | NR_115714  DQ207729 | Hungary | 12-MAR-2019 |
| *strain IAM 12077* | *B. thuringiensis* | NR_043403  D16281 | Japan | 18-MAR-2009 |
| *strain IAM 12605* | *B. cereus* | NR_115526  D16266 | Japan | 18-MAR-2009 |
| *strain ATCC 10792* | *B. thuringiensis* | NR_114581  AF290545 | USA | 02-OCT-2001 |
| *strain ATCC 14579* | *B. cereus* | NR_114582  AF290547 | USA | 02-OCT-2001 |
| *strain JCM 2152* | *B. cereus* | NR_113266  AB598737 | Japan | 26-NOV-2010 |
| *strain NBRC 101235* | *B. thuringiensis* | NR_112780  AB426479 | Japan | 05-MAR-2008 |
| *strain ATCC 14579* | *B. cereus* | NR_074540  AE016877 | France | 30-JAN-2014 |
| *strain NBRC 15305* | *B. cereus* | NR_112630  AB271745 | Japan | 05-SEP-2006 |
| *strain FZB42* | *B. velezensis* | NR_075005  CP000560 | Germany | 24-JUN-2019 |
| *strain 168* | *B. subtilis subsp. subtilis* | NR_102783  AL009126 | France | 08-FEB-2018 |
| *strain BGSC 3A28* | *B. subtilis subsp. inaquosorum* | NR_104873  HE582781 | Germany | 12-OCT-2011 |
| *strain DSM 10* | *B. subtilis* | NR_027552  AJ276351 | Germany | 08-JUL-2000 |
| *strain NBRC 13719* | *B. subtilis subsp. subtilis* | NR_112629  AB271744 | Japan | 05-SEP-2006 |
| *strain NBRC 101239* | *B. subtilis subsp. spizizenii* | NR_112686  AB325584 | Japan | 05-MAR-2009 |
| *strain JCM 1465* | *B. subtilis* | NR_113265  AB598736 | Japan | 02-JUN-2011 |
| *strain SBMP4* | *B. subtilis* | NR_118383  JQ424889 | India | 19-MAR-2012 |
| *strain ifo 15718* | *B. mojavensis* | NR_118290  JN585825 | China | 18-MAR-2014 |
| *strain NRRL B-23049* | *B. subtilis subsp. spizizenii* | NR_024931  AF074970 | USA | 04-JUN-1999 |
| *strain IFO15718* | *B. mojavensis* | NR_024693  AB021191 | USA | 26-APR-2000 |
| *strain NBRC 15718* | *B. mojavensis* | NR_112725  AB363735 | Japan | 12-OCT-2007 |
| *strain NBRC 100820* | *B. safensis* | NR_113945  AB681259 | USA | 28-JAN-2012 |
| *strain FO-36b* | *B. safensis* | NR_041794  AF234854 | USA | 19-SEP-2008 |
| *strain 41KF2b* | *B. altitudinis* | NR_042337  AJ831842 | India | 24-SEP-2008 |
| *strain NBRC 12092* | *B. pumilus* | NR_112637  AB271753 | Japan | 05-SEP-2006 |
| *strain ATCC 7061* | *B. pumilus* | NR_043242  AY876289 | USA | 11-AUG-2006 |
| *strain NBRC 15535* | *B. amyloliquefaciens* | NR_112685  AB325583 | Japan | 05-MAR-2009 |
| *strain NBRC 15535* | *B. amyloliquefaciens* | NR_041455  AB255669 | Japan | 05-OCT-2006 |
| *strain BCRC 11601* | *B. amyloliquefaciens* | NR_116022  EF433406 | Taiwan R.O.C. | 11-JUL-2008 |
| *strain MPA 1034* | *B. amyloliquefaciens* | NR_117946  HQ231913 | Germany | 03-NOV-2010 |
| *strain CBMB205* | *B. velezensis* | NZ_CP011937  CP011937 | Korea | 20-SEP-2018 |
| *strain IAM 12118* | *B. subtilis* | NR_112116  AB042061 | Japan | 14-JUN-2007 |
| *strain 168* | *B. subtilis subsp. subtilis* | NC_000964  AL009126 | France | 08-FEB-2018 |
| *strain MCCC 1A00732* | *B. nitratireducens* | NR_157732  KJ812430 | China | 27-JUL-2018 |
| *strain Bt407* | *B. thuringiensis* | NZ_CM000747  CM000747 | USA | 12-MAR-2015 |
| *strain HD 1* | *B. thuringiensis serovar kurstaki* | CP010005 | USA | 28-MAY-2015 |
